# Seamless workflow of hydroxy acid-modified metal oxide chromatography for rapid and sensitive phosphoproteomics sample preparation

**DOI:** 10.64898/2026.02.20.706508

**Authors:** Kazuya Tsumagari, Yasushi Ishihama, Koshi Imami

## Abstract

Phosphoproteomics by liquid chromatography/tandem mass spectrometry requires efficient phosphopeptide enrichment, but conventional workflows are often time-consuming and prone to sample loss, particularly at low input. Here, we present Rapid Hydroxy Acid–Modified Metal Oxide Chromatography (Rapid HAMMOC), a streamlined, TiO_2_-based enrichment workflow that features three key improvements. First, we optimized the TiO_2_ column loading conditions and found that alternative pH-buffering agents and organic solvents, such as sodium bicarbonate combined with ethyl acetate, outperformed the commonly used Tris-based buffer with isopropanol. Second, to minimize sample loss and manual handling in desalting, phosphopeptides eluted under basic conditions were directly loaded onto a dual-membrane StageTip composed of stacked strong anion exchange (SAX) and reversed-phase styrene–divinylbenzene (SDB) membranes (SAX-SDB StageTip). Third, the addition of lauryl maltose neopentyl glycol (LMNG), which is readily removed during desalting, suppressed nonspecific adsorption. Rapid HAMMOC provided markedly improved sensitivity, identifying approximately 5,000 class I phosphosites from 5 μg of K562 cell digests, with a median 7.9-fold increase in intensity compared to the original workflow. Rapid HAMMOC also identified, on average, approximately 8,000 class I phosphosites from as little as 0.5 μg input from HeLa, A549, and HCT116 cells. Furthermore, coupling Rapid HAMMOC with anti-puromycin immunoprecipitation enabled single-day profiling of nascent polypeptides from ultra-low input samples, yielding 2,310 high-confidence co-translational phosphosites. Beyond providing a practical enrichment workflow, this study offers broadly applicable insights that can be extended to other TiO_2_-based phosphoproteomic methods.

## Introduction

Understanding protein function is a critical objective in the post-genomic era. Phosphorylation of serine, threonine, and tyrosine is among the most common post-translational modifications and modulates the function of proteins, for example, by altering their activity, localization, and interactions ^1^. Abnormal phosphorylation is a hallmark of many diseases, including numerous cancers, Alzheimer’s disease, and diabetes ^1^. Liquid chromatography/tandem mass spectrometry (LC/MS/MS)-based proteomics is widely used to analyze phosphorylation status ^2^. For effective detection of phosphopeptides, enrichment from protein digests is a key step in sample preparation. Techniques such as metal oxide affinity chromatography (MOAC), using materials such as TiO_2_ (titania) and ZrO_2_ (zirconia) ^3,4^, and immobilized metal ion affinity chromatography (IMAC), which employs beads coordinated with metal ions such as Fe^3+^ and Ti^4+^^5,6^, are commonly used for this purpose. Sugiyama et al. previously reported hydroxy acid-modified metal oxide chromatography (HAMMOC) as a method for highly selective phosphopeptide enrichment, in which hydroxy acids such as lactic acid, β-hydroxypropanoic acid, and glycolic acid act as selectivity enhancers by preventing non-phosphorylated peptides from binding to metal oxides ^3^. The development of such enrichment techniques has been crucial for large-scale identification of phosphosites, facilitating system-wide understanding of the molecular mechanisms underlying phosphorylation-mediated biological processes and diseases ^7–11^.

A key recent advancement in proteomics is the development of platforms for analyzing data-independent acquisition (DIA) MS data ^12,13^, which enables faster and deeper analysis compared to conventional data-dependent acquisition (DDA) methods ^14,15^. With a current state-of-the-art LC/MS/MS system using DIA, it is now possible to profile up to 10,000 human proteins in half an hour ^15^ and to identify up to 7,400 proteins from as little as 250 pg of HeLa cell peptides ^14^. However, when targeting the phosphoproteome, sample amount often becomes a bottleneck due to the low stoichiometry of phosphorylation. To achieve sufficient depth in the phosphoproteome and cover key regulatory phosphosites, typical protocols for phosphopeptide enrichment require digests of 100–200 μg of protein per single shot ^13,16–18^, limiting their suitability for application to trace or high-cost samples. Sensitivity is not the only factor that limits biological research. A typical phosphopeptide enrichment method involves a low-throughput, time-consuming workflow, which includes two desalting steps—one after protein digestion and another after phosphopeptide enrichment—in addition to the enrichment step itself. This complexity represents a bottleneck that hinders the rapid processing of multiple samples and causes sample loss due to peptide adsorption onto tubes and pipette tips during transfer steps.

Streamlining the workflow not only improves operational efficiency but also enhances sensitivity by minimizing sample loss. Masuda et al. previously demonstrated an approximately 80-fold increase in sensitivity using an optimized phosphoproteomics sample preparation workflow applied to 10,000 HeLa cells ^19^. Proteins were extracted and digested using phase transfer surfactants (PTSs; i.e., sodium *N*-lauroylsarcosinate and sodium deoxycholate (SDC)) ^20,21^, and the resulting peptides were directly loaded onto a TiO_2_ column without desalting. Typically, peptide samples are loaded onto TiO_2_ columns under acidic conditions ^3,4,22^. However, anionic surfactants such as SDC precipitate under such conditions ^20^. To address this issue, acetonitrile (ACN) was added to solubilize the surfactants ^19^. This strategy was later revisited. Humphrey et al. improved the sensitivity further by replacing ACN with isopropanol (IPA) in the EasyPhos workflow, as IPA solubilizes SDC more efficiently ^17^. More recently, Oliinyk et al. systematically re-evaluated the entire workflow of EasyPhos for low-input samples and developed the more sensitive μPhos workflow ^23^. Furthermore, a recent preprint has described a phosphopeptide enrichment workflow for nanogram-scale samples, termed nanoPhos. The workflow employs solid-phase sample preparation, followed by automated phosphopeptide enrichment using Fe^3+^-NTA cartridges, thereby minimizing sample volumes and reducing opportunities for sample loss ^24^. Although these improvements have significantly increased the sensitivity and reduced the complexity of sample preparation, the desalting step for enriched phosphopeptides is unavoidable ^23,25^. In an IMAC-based workflow, Tsai *et al*. applied a tandem tip approach to the desalting step following phosphopeptide elution from the IMAC column, enabling phosphopeptides to be directly loaded onto a desalting column ^26^. Similarly, the nanoPhos workflow uses an Evosep system that enables high-throughput analysis while maintaining excellent sensitivity ^24,27^. However, when using TiO_2_ columns, phosphopeptides must be eluted under basic conditions, typically using reagents such as ammonia or piperidine ^17,19,25,28^, and subsequent desalting by reversed-phase chromatography requires acidic conditions. Therefore, the eluate must first be collected into a microtube, acidified (e.g., with trifluoroacetic acid; TFA), and then applied to the reversed-phase column. This step not only introduces the risk of sample loss due to peptide adsorption onto tubes and pipette tips, but also increases the need for manual handling, especially when processing multiple samples.

In this study, we revisited the HAMMOC method ^3^ and developed a streamlined, high-sensitivity phosphoproteomics sample preparation workflow, termed Rapid HAMMOC, using K562 cells as a model sample. To establish this method, we first compared various pH buffer components and organic solvents to determine the optimal conditions for phosphopeptide enrichment. We then developed a simple, efficient, and low-loss approach for phosphopeptide desalting, in which phosphopeptides eluted from the TiO_2_ column under basic conditions are directly applied to a desalting column without passing through an intermediate microtube. Furthermore, the addition of lauryl maltose neopentyl glycol (LMNG)—a surfactant that is readily removable during desalting— helped reduce nonspecific adsorption ^29^. We confirmed that the Rapid HAMMOC workflow is applicable not only to K562 cells but also to HeLa, A549, and HCT116 cells in the 0.5–20 μg range, enabling the identification of regulatory sites involved in intracellular signal transduction without activator treatment. Finally, we demonstrated that this workflow is applicable to ultra-low input samples, such as immunoprecipitates, exemplified by nascent polypeptide chains (NPCs) enriched using an anti-puromycin antibody, enabling the investigation of co-translational phosphorylation.

The importance of this study lies not only in presenting a practical workflow for phosphopeptide enrichment, but also in the fact that the concepts and findings described here are expected to be broadly applicable to other TiO_2_-based phosphopeptide enrichment methods.

## Materials and methods

### Materials

Roswell Park Memorial Institute 1640 medium (RPMI), Dulbecco’s modified Eagle’s medium (DMEM), SDC, sodium bicarbonate (NaBC), tris(2-carboxyethyl)phosphine (TCEP), 2-chloroacetamide (CAA), LysC, tris(hydroxymethyl)aminomethane (Tris), 4-(2-hydroxyethyl)-1-piperazineethanesulfonic acid (HEPES), DL-lactic acid, piperidine, ethyl acetate (EtOAc), LMNG, puromycin, bis(sulfosuccinimidyl) suberate disodium salt, and other reagents were purchased from FUJIFILM Wako (Osaka, Japan), unless otherwise specified. A phosphatase inhibitor cocktail was purchased from Merck (Darmstadt, Germany). Arginine- and lysine-free DMEM, a BCA assay kit, triethylammonium bicarbonate (TEAB), and Dynabeads™ Protein G magnetic beads were obtained from Thermo Fisher Scientific (Waltham, MA, USA). Stable isotope-labeled “heavy” arginine (L-[^13^C_6_,^15^N_4_], Arg10) and lysine (L-[^13^C_6_,^15^N_2_], Lys8) were obtained from Cambridge Isotope Laboratories (Tewksbury, MA, USA). Sequence-grade trypsin was obtained from Promega (Madison, WI, USA). TiO_2_ particles (Titansphere, 10 μm) and MonoSpin C18 columns were purchased from GL Sciences (Tokyo, Japan). C8, styrene-divinylbenzene (SDB), and strong anion exchange (SAX) membranes (Empore disks) were obtained from CDS Analytical (Oxford, PA, USA). Anti-puromycin monoclonal antibody (12D10) was obtained from Merck (Darmstadt, Germany).

### Cell culture

K562 cells were cultured in RPMI. HeLa, A549, and HCT116 cells were cultured in DMEM. All media were supplemented with 10% fetal bovine serum, 100 U/mL penicillin, and 100 μg/mL streptomycin. Cells were washed with ice-cold phosphate-buffered saline (PBS) prior to harvest and stored at −80 °C until use.

For the analysis of NPCs, HeLa cells were grown to approximately 60–70% confluence. The medium was then replaced with arginine- and lysine-free DMEM supplemented with 10% FBS, 10 μM puromycin, and heavy amino acids (0.398 mM arginine and 0.798 mM lysine), followed by incubation for 2 h.

### Protein extraction and digestion

Unless otherwise stated, cells were lysed in ice-cold 12 mM SDC buffer containing 100 mM NaBC, 10 mM TCEP, 40 mM CAA, and a phosphatase inhibitor cocktail ^20^. When an alternative pH-adjusting agent was used, its concentration was standardized to 100 mM. Lysates were immediately inactivated by heating at 95 °C for 5 min and then sonicated using a Bioruptor II (60 s on / 30 s off, 10 cycles; Sonicbio, Kanagawa, Japan). Protein concentration was determined using the BCA assay.

Proteins were digested overnight at 37 °C unless otherwise stated, with LysC and trypsin at a protein-to-enzyme ratio of 100:1 for each enzyme.

### Phosphopeptide enrichment

TiO_2_ columns were prepared by packing 0.5 mg of TiO_2_ particles onto C8 StageTips ^30^, constructed using a 10 μL pipette tip fitted with a 20-gauge C8 membrane frit. In the original HAMMOC workflow ^3^, the TiO_2_ column was equilibrated sequentially with 20 μL of 80% (v/v) ACN containing 0.1% (v/v) TFA (wash buffer), followed by 20 μL of 300 mg/mL lactic acid dissolved in wash buffer (enrichment buffer). The peptide sample, pre-purified using a MonoSpin C18 column or SDB StageTip and mixed with an equal volume of an enrichment buffer, was then loaded onto the TiO_2_ column. After sequential washing with 20 μL of enrichment buffer and 50 μL of wash buffer, phosphopeptides were eluted with 50 μL of 0.5% (v/v) piperidine (elution buffer). The eluate was acidified with TFA and purified using a SDB StageTip ^30^.

The Rapid HAMMOC workflow was established with the following modifications to the original protocol. Proteins were digested in SDC buffer containing 0.08% (v/v) LMNG ^29^ (typically, 20 μg of protein in 20 μL), and the resulting peptides were mixed with EtOAc and enrichment buffer at a sample:EtOAc:enrichment buffer ratio of 1:1:2. After agitation at 1,000 rpm for 20 min at room temperature, the mixture was loaded onto the TiO_2_ column. If precipitation was observed in the sample solution, the supernatant after centrifugation at 20,000 × *g* for 20 min was applied. Following sequential washes with enrichment buffer and wash buffer, phosphopeptides were directly eluted with elution buffer containing 0.02% LMNG onto a StageTip composed of 16-gauge SAX (top) and SDB (bottom) double membranes (SAX-SDB StageTip, see Fig. 2B) ^30^. Before sample loading, the SAX-SDB StageTip was pre-activated with 50 μL of wash buffer and equilibrated with 50 μL of elution buffer (no LMNG). After washing with 50 μL of 0.5% (v/v) TFA, phosphopeptides were eluted with 20– 30 μL of 40% (v/v) ACN containing 0.1% (v/v) TFA ^29^.

### Nascent polypeptide chain enrichment by pSNAP

Enrichment of NPCs using the pSNAP method was performed as previously described ^31^. Briefly, HeLa cells were lysed in a buffer containing 100 mM HEPES-NaOH (pH 7.5), 150 mM NaCl, 1% (v/v) Nonidet P-40 substitute, and protease and phosphatase inhibitor cocktails. Cell debris was removed by centrifugation at 16,000 × *g* for 30 min at 4°C. Protein concentration was determined using the BCA assay. Anti-puromycin antibodies (15 μg) were conjugated to 62 μL of Dynabeads™ Protein G magnetic beads per sample and then crosslinked using 5 mM bis(sulfosuccinimidyl) suberate disodium salt. The antibody-conjugated beads were incubated with 390 μg of proteins per sample for 1 h at 4°C with gentle rotation. The supernatant was discarded, and the beads were washed three times with ice-cold PBS supplemented with 850 mM NaCl. Nascent polypeptides were eluted with 20 μL of SDC buffer described above at 95°C for 5 min. The beads were then washed with an additional 10 μL of SDC buffer, which was combined with the eluate (final volume: 30 μL), and the resulting sample was subjected to the Rapid HAMMOC workflow. For digestion, 0.2 μg each of LysC and trypsin were added, and the mixture was incubated for 4 h at 37°C.

### Liquid chromatography/tandem mass spectrometry

All LC/MS/MS measurements were performed in DIA mode. In most experiments, a system comprising a Vanquish Neo UHPLC pump (Thermo Fisher Scientific) and an Orbitrap Eclipse mass spectrometer (Thermo Fisher Scientific) was used. The mobile phases consisted of (A) 0.1% (v/v) formic acid and (B) 80% (v/v) ACN containing 0.1% (v/v) formic acid.

In the method development experiments (Fig. 1), peptides were loaded onto a lab-made 15–18 cm fused-silica emitter packed with ReproSil-Pur C18-AQ (1.9 μm; Dr. Maisch, Ammerbuch, Germany) particles and separated by a linear gradient for 41 min (2.5–36% B over 30 min, 36–99% B over 1 min, and 99% B for 10 min) at the flow rate of 350 nL/min. All spectra were acquired using the Orbitrap analyzer. MS1 scans were acquired over an *m/z* range of 400–1,500 with a resolution of 120,000, maximum injection time of 45 ms, and automatic gain control (AGC) target of 300%. For subsequent MS/MS scans, the precursor range was set to *m/z* 450–1,250, and 80 scans were acquired with a 10 Th isolation window and additional 1 Th overlap. Higher-energy collisional dissociation (HCD) was applied with a normalized collision energy of 27. The scan range was set to *m/z* 110–1600, resolution to 15,000, injection time to 22 ms, AGC target to 1000%, and loop control to “All”.

**Figure 1.**
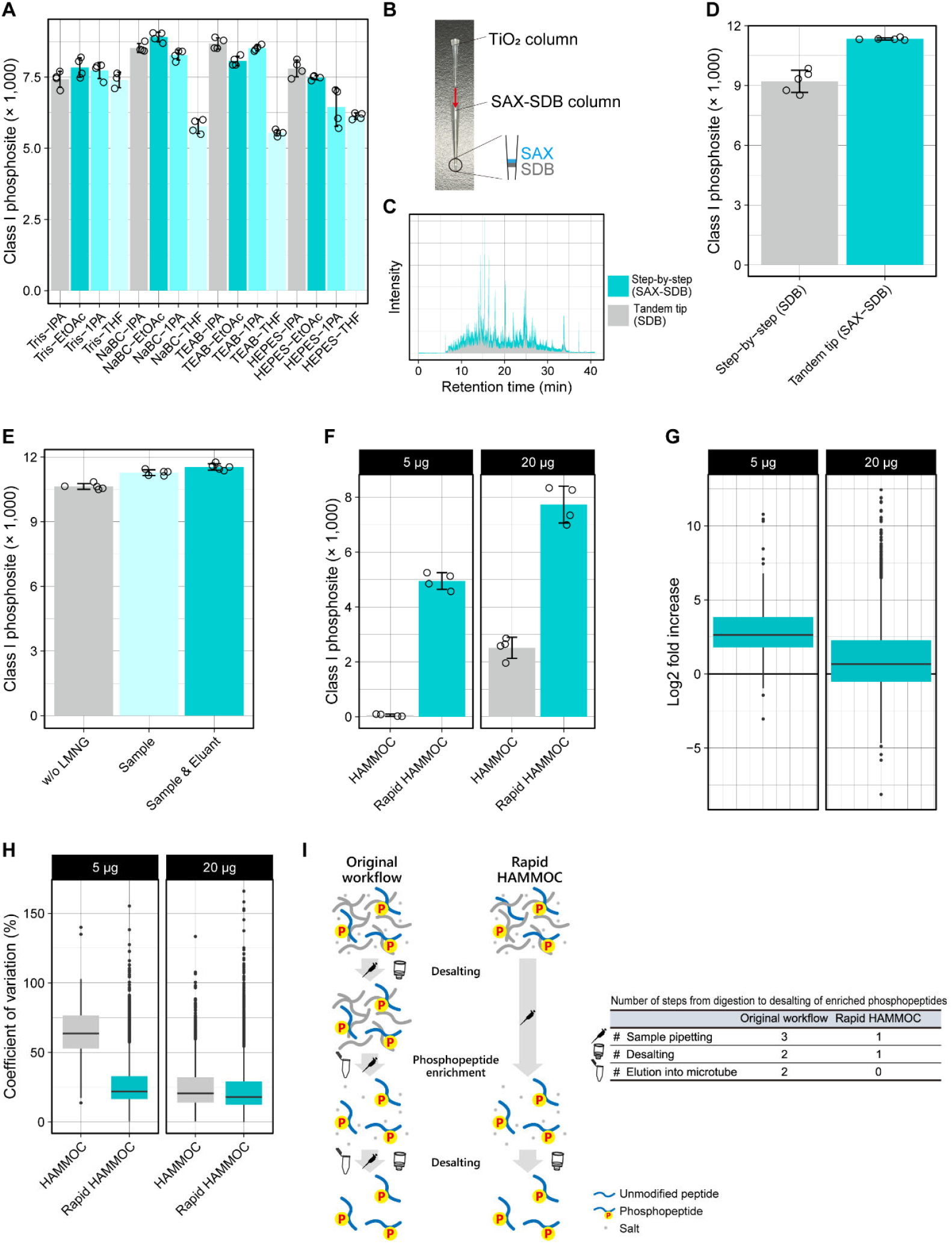
Optimization of the Rapid HAMMOC workflow. (**A**) Number of class I phosphosites identified with each sample loading condition. Four pH-buffering agents and four organic solvents are compared for all combinations. Each circle represents the number of phosphosites identified in a single sample. Bars indicate mean values, and error bars represent standard deviations (SDs). *N* = 4. NaBC, sodium bicarbonate; IPA, isopropanol; EtOAc, ethyl acetate; 1-PA, 1-propanol; THF, tetrahydrofuran. (**B**) Diagram showing the tandem tip-based desalting setup, connecting a TiO_2_ column to a SAX-SDB StageTip. (**C**) Representative total ion current chromatogram obtained with step-by-step SDB StageTip desalting or tandem tip desalting using SAX-SDB StageTip. (**D**) Number of class I phosphosites detected with step-by-step SDB StageTip desalting or tandem tip desalting using SAX-SDB StageTip. Each circle represents the number of phosphosites identified in a single sample. Bars indicate mean values, and error bars represent SDs. *N* = 5. (**E**) Number of class I phosphosites identified in samples without LMNG, with LMNG added to sample solutions, or with LMNG added to both sample and elution solutions. Each circle represents the number of phosphosites identified in a single sample. Bars indicate mean values, and error bars represent SDs. *N* = 5. (**F**) Number of class I phosphosites identified using the original HAMMOC workflow and Rapid HAMMOC workflow, with input amounts of 5 and 20 μg, respectively. Each circle represents the number of phosphosites identified in a single sample. Bars indicate mean values, and error bars represent SDs. *N* = 4. (**G**) Coefficients of variation (CVs) of class I phosphosite intensities quantified using the original HAMMOC workflow and Rapid HAMMOC workflow, with input amounts of 5 and 20 μg, respectively. Boxes represent the interquartile range (IQR), with the median shown as a horizontal line. Whiskers extend to 1.5 × IQR, and outliers are shown as individual points. (**H**) Log_2_ fold increases in class I phosphosite intensities observed with the Rapid HAMMOC workflow over the original HAMMOC workflow. Fold increases were calculated based on the mean values of replicates. Boxes represent the IQR, with the median shown as a horizontal line. Whiskers extend to 1.5 × IQR, and outliers are shown as individual points. (**I**) Schematic comparison of the original HAMMOC and Rapid HAMMOC workflows (Left). Number of sample pipetting, desalting, and elution steps into a microtube in the original HAMMOC workflow and Rapid HAMMOC workflow, respectively (Right).

In the method evaluation experiments (Fig. 2), the following modifications were applied: A 25 cm lab-made column was employed, and peptides were separated by a linear gradient for 85 min (2.5– 36% B over 70 min, 36–99% B over 5 min, and 99% B for 10 min) at the flow rate of 250 nL/min. For MS/MS scans, the resolution was set to 30,000, and maximum injection time to 54 ms.

**Figure 2.**
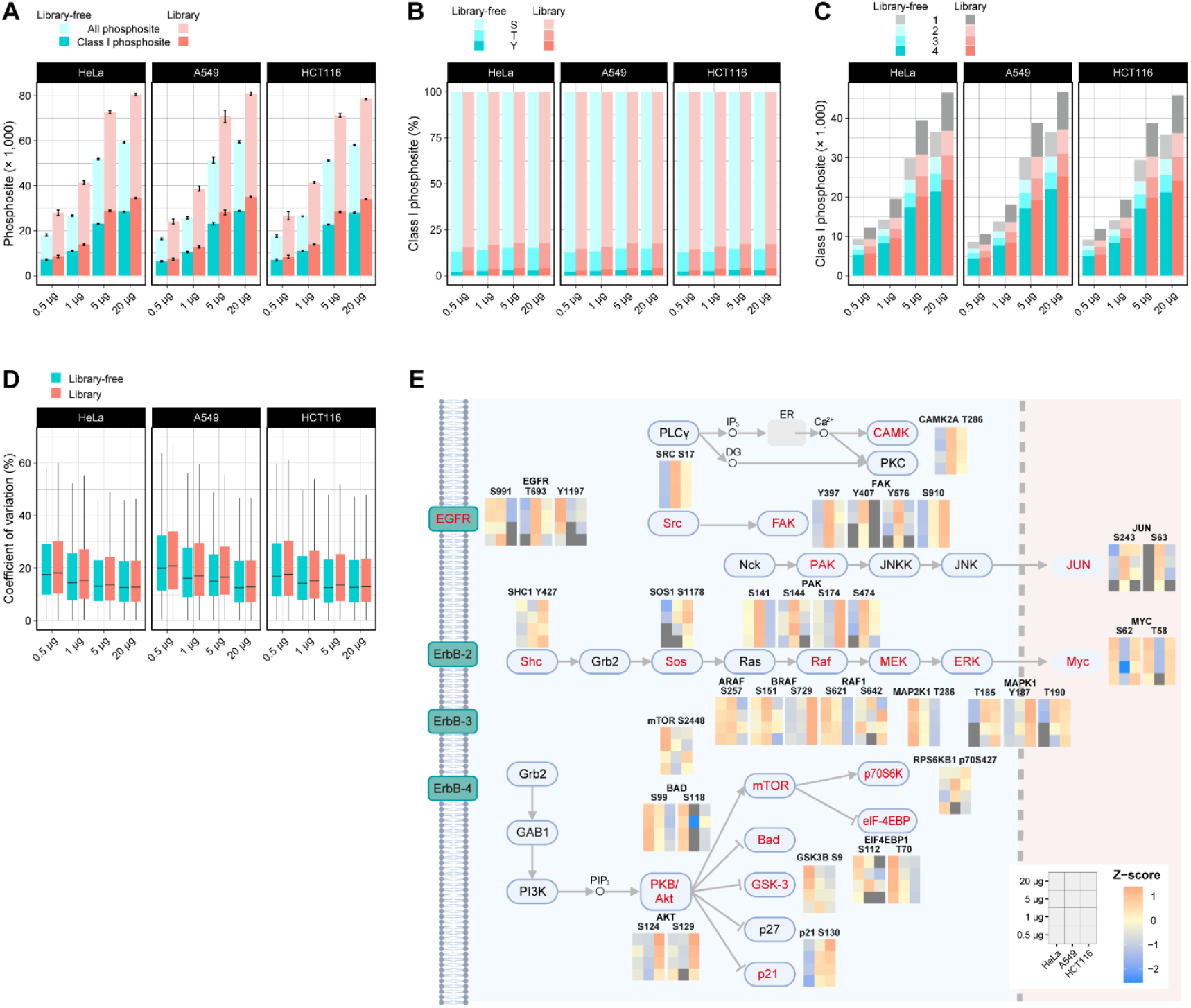
Evaluation of datasets generated using the Rapid HAMMOC workflow. (**A**) Number of identified phosphosites categorized by localization probability. Bars represent mean values, and error bars indicate standard deviations (SDs). *N* = 4. (**B**) Amino acid composition of class I phosphosites. (**C**) Number of class I phosphosites categorized by the number of valid quantifications across replicates. (**D**) Distribution of coefficients of variation (CVs) for class I phosphosites in each sample. Boxes represent the interquartile range (IQR), with the median shown as a horizontal line. Whiskers extend to 1.5 × IQR. Outliers are not shown. (**E**) Class I phosphosites mapped to the KEGG ErbB signaling pathway. Intensities are normalized as Z-scores within each input group, and mean values are shown for each sample. Only class I phosphosites that were quantified in at least one cell line with a 0.5 μg input and annotated as “functional sites” in PhosphoSitePlus are shown. (Created with BioRENDER.)

For measurement of phosphopeptides enriched from NPC samples (Fig. 3), an LC/MS/MS system comprising a Vanquish Neo UHPLC pump and an Orbitrap Astral mass spectrometer (Thermo Fisher Scientific) was used. Peptides were loaded on a 25 cm Aurora ULTIMATE column (AUR3-25075C18; IonOpticks, Collingwood, Australia) and separated by a linear gradient for 45 min (2.5–36% B over 35 min, 36–99% B over 2.5 min, and 99% B for 7.5 min) at the flow rate of 300 nL/min. MS1 scans were acquired using an Orbitrap analyzer over an *m/z* range of 400–1,500 with a resolution of 240,000, maximum injection time of 10 ms, and AGC target of 500%. Subsequent MS/MS scans were performed using an Astral analyzer. The precursor range was set to *m/z* 450–1,250, and 400 scans were acquired with a 2 Th isolation window and additional 1 Th overlap. The scan range was set to *m/z* 150-2,000, HCD normalized collision energy to 27, maximum injection time to 3.5 ms, AGC target to 500%, and loop control time to 0.6 s.

**Figure 3.**
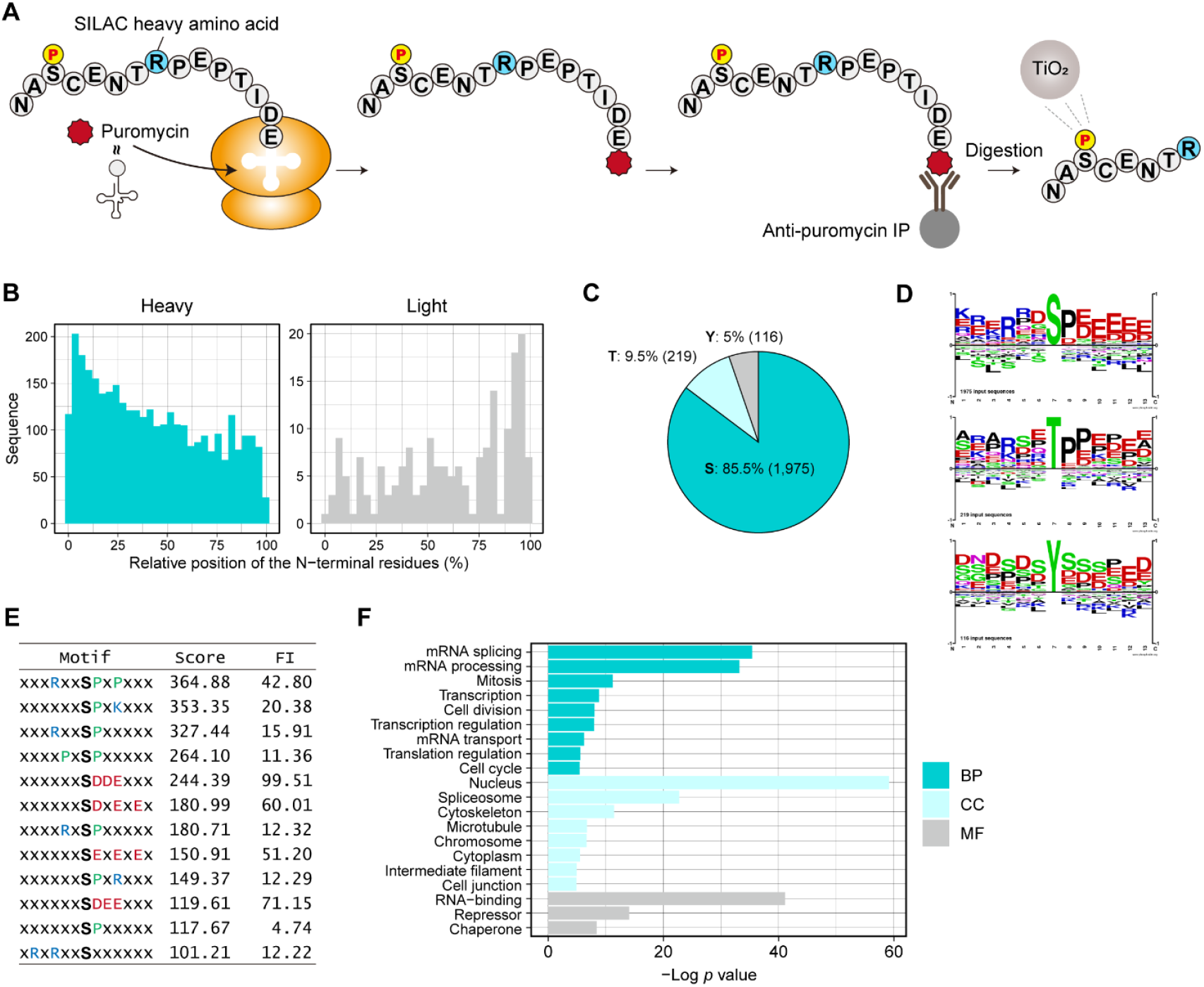
Profiling of co-translational phosphorylation on nascent polypeptide chains. (**A**) Workflow for profiling nascent polypeptide chains using pSNAP labeling followed by Rapid HAMMOC. Newly synthesized proteins are metabolically labeled with puromycin and SILAC heavy amino acids, which are supplied concurrently during pulse labeling. (**B**) Relative positions of the N-terminal residues of peptide sequences identified with SILAC heavy labeling within each protein (Left). Corresponding positions for peptide sequences detected exclusively with SILAC light labeling (Right). (**C**) Amino acid composition of class I phosphosites. (**D**) Sequence logos illustrating amino acid contexts surrounding co-translational phosphosites. (**E**) Overrepresented motifs identified from sequences flanking co-translational phosphosites. Only motifs with scores >100 are shown. No motifs meeting this threshold were detected for threonine or tyrosine. FI, fold increase. (**F**) Enrichment analysis of UniProt Keywords. An EASE threshold of 0.01 and a count threshold of 2 were applied. BP, biological process; CC, cellular component; MF, molecular function.

### MS data processing

MS raw files were processed using Spectronaut (v.19.7–20.1; Biognosys AG, Zurich, Switzerland). Throughout this study, the UniprotKB/SwissProt human database (April 2022), together with commonly observed contaminant proteins, was used for database searching. In the method development and evaluation experiments (Fig. 1 and 2), a DirectDIA (library-free) search was performed using the BGS phospho PTM workflow. Whole proteome data were analyzed using the BGS Factory Setting workflow. For library-based searches (Figs. 2 and 3), the BGS PTMs–Significant workflow was applied. In the analysis of co-translational phosphosites (Fig. 3), SILAC labeling was considered.

The PTM localization probability cutoff was set to 0 to report all identified phosphosites. Cysteine carbamidomethylation as a fixed modification, and oxidation on methionine, acetylation on the protein N-terminus, and phosphorylation on serine, threonine, and tyrosine as variable modifications were considered. Trypsin/P was set as the cleavage enzyme with a maximum of two missed cleavages. The mass tolerance was set to dynamic in both MS1 and MS2 with a correction factor of 1. A Q value of 1% against mutated decoys was applied to filter identifications with a false-discovery rate of 0.01 at both the peptide and protein group levels. Quantification was performed using the automatic setting. In the method development experiments (Fig. 1), the cross-run normalization feature was disabled.

### Spectral library construction

Spectral libraries were constructed by using Spectronaut with the database, fixed and variable modifications, and cleavage specificity settings described above. In the method evaluation experiment (Fig. 2), two spectral libraries were used based on the sample size-compatible spectral library concept ^32^. The “small library” was generated from data acquired using 0.5–20 μg of input from HeLa, A549, and HCT116 cells. The “large-scale library” was generated by combining all data generated in the method evaluation experiment with additional data acquired under the same LC/MS/MS conditions, using 100 μg of digested materials from the same cell lines. This supplementary dataset was prepared using the original HAMMOC protocol ^3,28^. The small library was used for samples with 0.5–5 μg input, while the large library was used for samples with 20 μg input.

In the analysis of phosphosites on NPCs (Fig. 3), a project-specific spectral library was initially generated from the acquired raw data. SILAC light and heavy (Arg10 and Lys8) labels were assigned to channels 1 and 2, respectively. Additionally, a HeLa cell-specific phosphoproteome spectral library was generated from measurement data acquired using input amounts of 0.5, 1, and 5 μg (Fig. 2) ^32^. Based on the spectral information for the light (unlabeled) ions, corresponding spectra for SILAC heavy-labeled ions were computationally predicted and incorporated. These two spectral libraries were concatenated and used for database searching.

### Bioinformatics and statistical analysis

In method development and evaluation experiments (Figs. 1 and 2), PTM site reports in BGS Factory report format were exported for downstream data analysis and interpretation. Rows were filtered to retain those with quantity value >1. For phosphosites that were redundantly reported due to mapping to multiple proteins within a protein group, only those assigned to the primary protein accession in the PG.ProteinGroups column were considered. Non-log-transformed intensities were used to calculate the coefficients of variation. In the analysis of co-translational phosphosites (Fig. 3), precursor reports in BGS Factory report format were exported and converted to site-level matrices using a Peptide Collapse plugin in Perseus (v.1.6.14) ^13,33^, with modifications: to output all detected phosphosites, missing values (NA, 0, 1) were replaced with the minimal quantity.

Data analysis and visualization were performed using the R framework (v.4.4.2) with the basic functions and the ggplot2 (v.3.3.6) packages and Perseus (v.1.6.14) ^33^. KEGG pathway mapping and visualization were performed by enrichment analysis in DAVID ^34^, BioRENDER, R, and Illustrator (Adobe, San Jose, CA, USA). UniProt Keywords enrichment analysis was performed using DAVID ^34^. Sequence logos were generated using PhosphoSitePlus (https://www.phosphosite.org/). Motif analysis was performed using motif-X 2.0 (https://proteome-x.com/motif-x) ^35^.

### Experimental Design and Statistical Rationale

In method optimization experiments (Fig. 1), differences between conditions were evaluated using identical cell lysates in each experiment, and 4–5 replicate samples were prepared per condition. In the pH-buffering agent comparison, cells were first lysed in buffer lacking any pH-buffering agent, then equally divided, and pH-buffering agents were added separately, ensuring equal input across conditions. Unless otherwise specified, NaBC was used as the pH-controlling agent in the digestion buffer. SAX-SDB StageTips were consistently used for phosphopeptide desalting throughout method development, not only in comparative experiments involving SDB and SAX-SDB StageTips. Unless otherwise specified, raw files were searched within individual conditions. When calculating fold changes in intensities between different conditions, the samples were searched together in a single database search. In method evaluation experiments (Fig. 2), database search was performed within each input group to enable comparison between cell lines. In the analysis of co-translational phosphosites (Fig. 3), three pSNAP sample replicates were prepared from the same HeLa cell pellet and subsequent phosphopeptide enrichment was applied to each replicate.

## Results and discussion

### Optimizing the HAMMOC workflow to improve experimental efficiency and sensitivity

Based on the HAMMOC method ^3^, we optimized the sample preparation workflow in phosphoproteomics with the aim of improving experimental efficiency and sensitivity, using digests of 20 μg of protein extracted from K562 cells as the input per sample.

For the enrichment of phosphopeptides from small sample amounts, recent studies have shown that omitting the desalting step after digestion and directly applying the digest to a TiO_2_ column is a promising strategy for improving sensitivity by minimizing sample loss due to adsorption onto tips or columns ^17,19,23,25^. In particular, following digestion in a solution containing SDC, a workflow dissolving SDC under acidic conditions using IPA and directly loading the digest onto a TiO_2_ column has become widely adopted ^17,23,25^. However, it remains unclear whether IPA is the optimal organic solvent in terms of phosphosite identification efficiency, as no detailed comparative studies have been reported. Similarly, Tris-HCl, which is commonly used for pH adjustment of digestion buffers, has not been compared with other pH-buffering agents. Therefore, first, we conducted a comparative evaluation of pH buffer components and organic solvents (Fig. 1A). K562 cell proteins were digested in SDC-based buffers adjusted to pH 8.5–9.0 using Tris-HCl, sodium bicarbonate (NaBC), triethylammonium bicarbonate (TEAB), or HEPES-NaOH. Subsequently, IPA, EtOAc, 1-propanol (1-PA), or tetrahydrofuran (THF) was added, followed by lactic acid as a selectivity enhancer (see the Materials and Methods section for details) ^3^. We evaluated the results based on the number of detected phosphosites with a localization probability >0.75, which is commonly accepted as the criterion for confident phosphosites, class I phosphosites, in the proteomics community. The combination of NaBC and EtOAc yielded the highest number of identified class I phosphosites, closely followed by the combination of TEAB and IPA. Notably, many of the conditions with NaBC or TEAB, outperformed the widely used Tris-based digestion buffer, underscoring the significance of these findings. There were no substantial differences in digestion efficiency or the total number of identified precursors among the different digestion buffers.

In phosphopeptide enrichment workflows using TiO_2_ columns, phosphopeptides are typically eluted using a basic solution. However, since the downstream desalting step requires an acidic condition, phosphopeptides must first be eluted into a microtube, where the eluate is either acidified or subjected to buffer exchange prior to desalting ^3,19,23,25^. To address this limitation, we sought to develop a method to directly trap phosphopeptides eluted from the TiO_2_ column onto the desalting column using a basic solution. Specifically, we utilized a dual-membrane StageTip composed of stacked strong anion exchange (SAX) and reversed-phase styrene–divinylbenzene (SDB) membranes (SAX-SDB StageTip) ^30^. This approach builds upon the concept of tandem-tip workflows ^26^ and enables direct elution of phosphopeptides from the TiO_2_ column into the SAX-SDB StageTip, thereby minimizing sample loss (Fig. 1B). This strategy led to a clear improvement in sensitivity (Fig. 1C), as evidenced by the increased number of identified class I phosphosites (Fig. 1D), and also enhanced experimental efficiency, as reflected by the reduced number of sample pipetting steps in the overall workflow.

The use of MS-compatible surfactants has been demonstrated to reduce peptide adsorption onto tubes and pipette tips ^25,29,36^. However, n-dodecyl-β-D-maltoside (DDM), a surfactant that has recently been frequently employed for this purpose, co-elutes with peptides during LC/MS/MS analysis ^29^, potentially interfering with phosphopeptide detection. Here, we tested the use of LMNG, which is readily removed during desalting ^29^. The addition of LMNG during sample preparation resulted in a trend toward increased identification of class I phosphosites (Fig. 1E). Furthermore, the inclusion of LMNG in the elution buffer used for phosphopeptide elution onto the SAX-SDB StageTip resulted in a further increase in the number of identified class I phosphosites.

We then integrated the three optimized components—(1) digestion conditions and SDC-solubilization organic solvent selection, (2) simplification of the desalting step using the SAX-SDB StageTip, and (3) reduction of adsorption loss through LMNG supplementation—into a unified Rapid HAMMOC workflow, and compared it to the original workflow. This optimized workflow was applied to 5 μg and 20 μg of protein extracted from K562 cells. Notably, using the Rapid HAMMOC workflow, approximately 5,000 class I phosphosites were identified on average from just 5 μg of input, whereas fewer than 100 sites were detected using the original protocol (Fig. 1F). With 20 μg of input, the number of identified sites increased by approximately 3.1-fold. Sensitivity improvements were further supported by increased intensities (Fig. 1G), with a median 7.9-fold increase in the 5 μg input samples and a 2.1-fold increase in the 20 μg input samples. The median coefficients of variation (CVs) improved from 63.6% to 21.9% with 5 μg input, and from 20.6% to 17.9% with 20 μg input (Fig. 1H). Together, these results clearly demonstrate the effectiveness of the Rapid HAMMOC workflow in terms of both sensitivity and reproducibility.

The improvements are summarized in Fig. 1I. The Rapid HAMMOC workflow eliminates multiple elution steps into a microtube and requires only one desalting step. Moreover, only a single sample pipetting step—loading onto the TiO_2_ column—is required between digestion and the desalting of enriched phosphopeptides. This streamlined procedure minimizes manual handling, enabling rapid and efficient phosphoproteome sample preparation compared to conventional workflows.

Importantly, the benefits demonstrated here should also extend to other TiO_2_-based phosphopeptide enrichment strategies, highlighting the potentially broad applicability of our approach in the phosphoproteomics field.

### Evaluating the practical utility of the Rapid HAMMOC workflow in HeLa, A549, and HCT116 cells

We applied the Rapid HAMMOC workflow to a dilution series of 20, 5, 1, and as little as 0.5 μg of input material prepared from HeLa, A549, and HCT116 cells. For each condition, both library-free and library-based searches were performed. Different spectral libraries were used depending on the input amount, following the strategy described previously (see the Materials and Methods section for details) ^32^.

Overall, the numbers of identified phosphosites were comparable across the three cell lines (Fig. 2A). Using the library-free search, we identified an average of 6,849, 10,858, 22,970, and 28,386 class I phosphosites from 0.5, 1, 5, and 20 μg of input, respectively. Similarly, with the library-based search, 8,097, 13,509, 28,502, and up to 34,525 class I phosphosites were identified on average.

Compared to serine and threonine phosphorylation, which predominated in all conditions, tyrosine phosphorylation was detected at lower frequencies, likely due to its low stoichiometry, despite its known regulatory importance (Fig. 2B) ^37^. In library-free searches, class I tyrosine phosphosites accounted for, on average, 1.9%, 2.4%, 3.0%, and 2.8% of all class I phosphosites in the 0.5, 1, 5, and 20 μg input samples, respectively, highlighting the challenge of detecting low-stoichiometric tyrosine phosphosites in low-input samples. The use of library-based searches improved detection: the proportions of tyrosine phosphosites increased to 2.8%, 3.6%, 4.0%, and 4.0%, respectively, in the same samples.

When label-free quantification is used, missing values can pose problems and complicate downstream statistical analysis, although they are generally less frequent in DIA compared to DDA. With library-free search, more than 50% of class I phosphosites were quantified across all four replicates, and approximately 70% were quantified in at least three replicates (Fig. 2C). In the library search, the proportion of frequently quantified sites was slightly lower: on average, 60–66% of sites were quantified at least three times. This may be attributed to the inclusion of more low-intensity precursors near the limit of quantification in library search. In the library-free search results, the median CVs were on average 18.1%, 15.0%, 13.6%, and 12.7% in the 0.5, 1, 5, and 20 μg input samples, respectively, demonstrating good reproducibility (Fig. 2D). Median CVs tended to be slightly higher in the library-based search results: 18.9%, 15.9%, 14.7%, 12.9%, respectively.

These findings show that the Rapid HAMMOC workflow can generate practical datasets even from small input amounts such as 0.5 μg. Indeed, when we mapped the data onto the KEGG ErbB signaling pathway—a representative intracellular signaling cascade—we observed that functional phosphosites annotated in PhosphoSitePlus were broadly represented, from upstream EGFR, through the MAP kinase cascade, to downstream MYC (Fig. 2E).

In summary, Rapid HAMMOC enables sensitive and reproducible phosphoproteomic profiling from low-input samples. To benchmark its performance, we compared it with other low-input phosphopeptide enrichment methods. For example, the SOP-Phos method reported the identification of an average of 3,436, 5,369, 8,602, and 12,896 phosphopeptides from 0.5, 1, 2.5, and 5 μg of PC9 cell digests, respectively, using an Orbitrap Fusion Lumos Tribrid mass spectrometer with library-free search ^25^. Similarly, the μPhos method identified approximately 17,000 class I phosphosites from 20 μg of HeLa digests using a timsTOF Ultra mass spectrometer and a library-free search. More than 12,000 sites were still detected with half the input, and with one-quarter, only around 5,000 fewer sites were identified. Even with as little as 1 μg of input, up to 6,200 class I phosphosites were detected ^23^. Compared to these high-sensitivity phosphopeptide enrichment strategies, Rapid HAMMOC demonstrated comparable sensitivity. Moreover, it offers a streamlined and robust workflow without the need for DDM pre-coating of microtubes ^25^ and with a simplified desalting procedure. These improvements not only underscore the value of the Rapid HAMMOC workflow itself, but again, importantly, offer insights that should be applicable to improve the experimental efficiency and sensitivity of other TiO_2_-based phosphopeptide enrichment methods.

### Exploring co-translational phosphosites mapped on nascent polypeptide chains

Modifications such as protein N-terminal acetylation and myristoylation are introduced co-translationally during the process of protein synthesis ^38^. Phosphorylation can also occur at this stage, and for several protein kinases—including PKA, GSK3β, and DYRK1A—phosphorylation during biosynthesis has been shown to be essential for maintaining their stability and functional activity ^39– 43^. However, conventional phosphoproteomics workflows are unable to distinguish co-translational phosphorylation events from those that occur post-translationally. Our group previously developed the pSNAP method for proteome-wide profiling of nascent polypeptide chains (NPCs), which employs affinity enrichment using an anti-puromycin antibody ^31^. A key strength of the pSNAP approach is that NPCs are metabolically pulse-labeled with both puromycin and SILAC heavy amino acids during protein synthesis, enabling a clear distinction between genuine NPCs and nonspecific contaminants. The amount of NPCs obtained via immunoprecipitation is typically less than 1 μg, making it challenging to apply conventional phosphopeptide enrichment protocols. To address these challenges, we finally investigated co-translational phosphosites in HeLa cells by integrating pSNAP with the Rapid HAMMOC method (Fig. 3A). Notably, sample preparation steps —from NPC enrichment to phosphopeptide enrichment—were completed within a single day by employing a shortened digestion time of 4 h ^23,25^.

Using the state-of-the-art Orbitrap Astral mass spectrometer, we identified a total of 8,770 heavy-labeled precursors from measurements of three sample replicates, corresponding to 3,082 phosphosites mapped across 3,409 peptide sequences derived from 1,414 proteins. Consistent with the directional nature of ribosomal protein synthesis, these heavy-labeled phosphopeptides exhibited a clear bias toward the N-termini of the corresponding proteins (Fig. 3B). In contrast, phosphopeptides that were detected exclusively as unlabeled (light) forms showed no such trend. Among them, 2,310 sites exhibited high localization probability (> 0.75), categorized as class I phosphosites, which were used for further analysis. Of these, 85.5% were serine residues, followed by threonine (9.5%) and tyrosine (5%) (Fig. 3C). Notably, two known co-translational phosphosites, GSK3A Y279 and DYRK1A Y321, were included in the identified class I phosphosites ^40–43^.

Sequence logos illustrating the amino acid context surrounding the phosphosites are shown in Fig. 3D. In general, for both serine and threonine phosphosites, basic residues were predominantly enriched on the N-terminal side, whereas acidic residues were more frequently observed on the C-terminal side. Motif enrichment analysis revealed several significantly overrepresented sequence patterns consistent with known kinase recognition motifs (Fig. 3E). Proline-directed motifs centered on SP, commonly associated with MAPK and CDK family kinases, were prominently enriched. In particular, proline-directed motifs with a basic residue at the +3 position, such as SPxK and SPxR, aligned with the substrate preferences of CDK family kinases, while a motif containing two prolines at the –2 and +1 positions, namely PXSP, was consistent with the recognition motifs of MAPK family kinases ^44^. Acidophilic motifs containing clusters of D/E residues, such as SDDE, SDxExE, SExExE, and SDEE, were also strongly overrepresented, consistent with the preference of CK2 ^45^. Additionally, a basophilic motif containing multiple basic residues, such as RxRxxS, was consistent with the substrate specificity of AGC family kinases, specifically matching the canonical recognition motif of AKT ^44^. Enrichment analysis of UniProt Keywords revealed involvement of co-translational phosphorylation in various classes of proteins in terms of biological process, molecular function, and intracellular localization (Fig. 3F).

Despite its biological relevance, the functional role of co-translational phosphorylation has remained poorly characterized, primarily due to technical limitations. We identified more than 2,000 high-confidence co-translational phosphosites in NPCs enriched via anti-puromycin immunoprecipitation, highlighting the exceptional sensitivity of the Rapid HAMMOC method. Furthermore, the use of short digestion enabled the preparation of phosphoproteome samples to be completed within a single day, underscoring the efficiency of our workflow. Our findings revealed that co-translational phosphorylation targets not only protein kinases, but also a broad spectrum of other proteins, suggesting that it functions as a general regulatory mechanism involved in protein stability and folding. Continued investigation into co-translational phosphorylation may uncover previously unrecognized functions of this regulatory modification.

## Conclusions

In this study, we developed a streamlined and sensitive phosphoproteomics sample preparation workflow, termed Rapid HAMMOC, that is especially suitable for low-input samples. By using Rapid HAMMOC, we achieved a dramatic increase in the number of identified class I phosphosites and an improvement in quantitative reproducibility. We would like to emphasize that the ideas and findings described here should also be applicable to other TiO_2_-based phosphopeptide enrichment methods, and thus a broad impact on phosphoproteomics is anticipated. The utility of this method was demonstrated by applying it to three cell lines across an input range of 0.5 to 20 μg, enabling the identification of over 8,000 class I phosphosites even from low-input (0.5 μg) samples, and allowing profiling of functional phosphosites in the ErbB signaling pathway of these cell lines in the steady state. Furthermore, by applying Rapid HAMMOC to anti-puromycin antibody immunoprecipitation samples from HeLa cells labeled with puromycin on NPCs, we were able to identify co-translational phosphosites on nascent chains, demonstrating the high sensitivity of this method. In summary, Rapid HAMMOC can be easily implemented in a proteomics laboratory as an extended workflow of routine phosphoproteome sample preparation, contributing to sensitive and reproducible analysis.

## Acknowledgments

This work was supported by Japan Science and Technology Agency (JST) ERATO Arita Lipidome Atlas Project (JPMJER2101 to KI), JST FOREST (JPMJFR214L to KI), Japan Society for the Promotion of Science (JSPS) Grant-in-Aid for Early-Career Scientists (23K14169 to KT), JSPS Grant-in-Aid for Scientific Research (20H03241, 20H04844, 21H05720 to KI; 23H04924 to KI and YI; 25K22526 to YI), and the RIKEN Special Postdoctoral Researcher program (to KT). We thank Kanako Igarashi (RIKEN) for technical support. ErbB pathway map was created in BioRender (Tsumagari, K. (2026) https://BioRender.com/8170nk9).

## Notes

The authors declare no competing financial interest.

## Notes

### Competing Interest Statement

The authors have declared no competing interest.

